# The mitochondrial Na^+^/Ca^2+^ exchanger NCLX is implied in the activation of hypoxia-inducible factors

**DOI:** 10.1101/2024.07.12.603081

**Authors:** Carmen Choya-Foces, Pablo Hernansanz-Agustín, Elisa Navarro, Manuela G. López, Javier Egea, Antonio Martínez-Ruiz

## Abstract

Eukaryotic cells and organisms depend on oxygen for basic living functions, and they display a panoply of adaptations to situations in which oxygen availability is diminished (hypoxia). A number of these responses in animals are mediated by changes in gene expression programs directed by hypoxia-inducible factors (HIFs), whose main mechanism of stabilization and functional activation in response to decreased cytosolic oxygen concentration was elucidated two decades ago. Human acute responses to hypoxia have been known for decades, although their precise molecular mechanism for oxygen sensing is not fully understood. It is already known that a redox component, linked with reactive oxygen species (ROS) production of mitochondrial origin, is implied in these responses. We have recently described a mechanism by which the mitochondrial sodium/calcium exchanger, NCLX, participates in mitochondrial electron transport chain regulation and ROS production in response to acute hypoxia.

Here we show that NCLX is also implied in the response to hypoxia mediated by the HIFs. By using a NCLX inhibitor and interference RNA we show that NCLX activity is necessary for HIF-α subunits stabilization in hypoxia and for HIF-1-dependent transcriptional activity. We also show that hypoxic mitochondrial ROS production is not required for HIF-1α stabilization under all circumstances, suggesting that other mechanism(s) could be operating in the NCLX-mediated response to hypoxia that operates through HIF-α stabilization. This finding provides a link between acute and medium-term responses to hypoxia, reinforcing a central role of mitochondrial cell signalling in the response to hypoxia.

## Introduction

Most living organisms depend on oxygen for their vital functions, especially as an electron acceptor in the respiration process, which in eukaryotic cells takes place in mitochondria. Thus, a variety of adaptations have been selected to respond to hypoxia, i.e. the situation in which less oxygen is available. In humans and some mammals, acute responses to hypoxia such as hyperventilation or pulmonary vasoconstriction depend on specialized cells that transduce reduced oxygen availability in the blood or in their environment in signals that elicit the responses. The precise molecular identity of the oxygen sensors involved is not fully known, but it is clear that changes in redox state and in reactive oxygen species (ROS) production are involved in the signal initiation and/or transduction, including a main role of mitochondrial complex I (Fernández-Agüera et al., 2015; Hernansanz-Agustín et al., 2017).

We have recently shown a molecular mechanism by which mitochondria produce reactive oxygen species (ROS) in an acute response to hypoxia, with a central role of mitochondrial Na^+^ import mediated by the mitochondrial Na^+^/Ca^2+^ exchanger, NCLX (Hernansanz-Agustín et al., 2020). Mitochondrial complex I undergoes the active/deactive (A/D) transition when oxygen levels are decreased, which involves the halt of its NADH:CoQ oxidoreductase and H^+^ pumping activities, and leads to a slight acidification of the mitochondrial matrix (Hernansanz-Agustín et al., 2017). This acidification partially dissolves the mitochondrial calcium phosphate precipitates and releases soluble Ca^2+^ into the matrix which, in turn, activates the mitochondrial Na^+^/Ca^2+^ exchanger (NCLX). NCLX extrudes Ca^2+^ in exchange of Na^+^ which, once inside the matrix, interacts with the phospholipids in the inner leaflet of the inner mitochondrial membrane, uncoupling the Q-cycle in complex III. Ultimately, complex III produces superoxide at the level of its Qo site during the first minutes of hypoxia (Hernansanz-Agustín et al., 2020). Interestingly, this work showed that inhibition of NCLX either pharmacologically or genetically did not alter respiration rate.

On the other hand, medium-term genetic adaptations to hypoxia rely mostly on a family of transcription factors, namely the hypoxia-inducible factors (HIFs), which are activated after several hours of hypoxia. HIFs are composed of an α subunit and a β subunit. The latter is constitutively expressed, while the former is continuously synthetized and degraded and is only stable under conditions of low oxygen, high levels of hydrogen peroxide (H_2_O_2_) or other metabolic challenges (Bruick & McKnight, 2001; Epstein et al., 2001; Gerald et al., 2004; Ivan et al., 2001; Selak et al., 2005). During normoxia, HIF-α subunits become hydroxylated by specific HIF prolyl-hydroxylases (PHDs) that target them for ubiquitination by the Von Hippel Lindau (VHL) factor and degradation by the 26S proteasome. However, under hypoxia, PHDs are not able to perform the hydroxylation of HIF-α anymore and the subunits stabilize, allowing their dimerization with the β subunit and the initiation of the hypoxic transcriptional programme (Kaelin & Ratcliffe, 2008).

Although PHDs have been demonstrated to depend on O_2_ to perform the hydroxylation reaction and are mainly located in the cytosol and the nucleus, HIF-α stabilization has also been observed to depend on ROS production by mitochondria (Brunelle et al., 2005; Chandel et al., 1998; Guzy et al., 2005; Mansfield et al., 2005; Fuhrmann & Brüne, 2017). Mitochondrial ROS increase during the first minutes of hypoxia (Hernansanz-Agustín et al., 2014, 2021) involving respiratory complexes I and III (Brunelle et al., 2005; Chandel et al., 1998, 2000; Guzy et al., 2005; Hernansanz-Agustín et al., 2017; Mansfield et al., 2005). Since the systems used to study the involvement of mitochondrial ROS on HIF-α stabilization implied the ablation or inhibition of either complex I or complex III, they also compromised the respiratory machinery, thus modifying the O_2_ gradients in the cell (Hagen et al., 2003), among other metabolic parameters (Martínez-Reyes et al., 2016). This made impossible to differentiate whether the prevention of HIF-α stabilization was due to the modification of such gradients or the blockade of ROS production (Kaelin & Ratcliffe, 2008).

Here we show that inhibition or interference of NCLX (without altering the mitochondrial electron transport chain complexes stability or respiratory function) abolishes the response to hypoxia mediated by HIF-α subunit stabilization and transcriptional activity in different cell types, linking the acute and medium-term response to hypoxia. However, particular cells do maintain the HIF response to hypoxia after NCLX inhibition or ablation, while their hypoxic ROS production is still dependent on NCLX activity. We found that this occurs due to higher basal cytosolic oxidative status, suggesting that the redox role of mitochondria in HIF-α stabilization during hypoxia may be masked by alterations on the basal oxidant/antioxidant system.

## Materials and Methods

### Cell culture, brain slices, treatments and hypoxic exposure

Cells were routinely maintained in cell culture incubators (95% air, 5% CO_2_ in gas phase, 37 °C). Bovine aortic endothelial cells (BAECs) were isolated as described (Navarro-Antolín et al., 2000) and cultured in RPMI 1640 supplemented with 15% heat-inactivated fetal bovine serum (FBS), 100 U/mL penicillin and 100 μg/mL streptomycin. Human umbilical vein endothelial cells (HUVECs) were isolated as described (Muñoz et al., 1996) and cultured in Medium 199 supplemented with 20% heat-inactivated FBS, 16 U/mL heparin, 100 mg/L ECGF (endothelial cell growth factor), 20 mM HEPES (4-(2-hydroxyethyl)piperazine-1-ethanesulfonic acid), 100 U/mL penicillin and 100 μg/mL streptomycin. BAECs were used between passages 3 and 9 and HUVECs between passages 3 and 7. Endothelial morphology was assessed by visual inspection and by expression of endothelial nitric oxide synthase (eNOS). Immortalized mouse hippocampal cells (HT22 cell line) HT22 cells were cultured in DMEM, supplemented with 10 % heat-inactivated FBS, 20 mM HEPES, 100 units/mL penicillin, and 100 μg/mL streptomycin.

Mouse embryonic fibroblasts (MEFs) obtained from C57BL/6N mice harbouring LoxP regions in the *Slc8b1* gene (*Slc8b1* fl/fl) were kindly provided by Dr. John Elrod (Luongo et al., 2017). Mouse adult fibroblasts (MAFs) isolated from adult (20 weeks old) C57BL/6N mouse ears were kindly provided by Dr. José Antonio Enríquez. Both cell types were cultured in DMEM supplemented with 10% heat-inactivated FBS, 100 U/mL penicillin and 100 μg/mL streptomycin.

MEFs were immortalized by retrovirus infection. 293T cells were transfected with 12 ug pCL-Ampho and 12 ug pBABE-SV40-puro using Lipofectamine RNA iMAX (Invitrogen). Two days later, 1/2 dilution of 293T cells media and 8 ug/mL of polybrene were added on MEFs culture for 4 hours. Then, the infection was repeated, but adding 4 ug/mL of polybrene overnight. Next day, the first infection was repeated and, upon 4 hours, MEFs were passaged with 1/1000 of puromycin. Acute ablation in *Slc8b1* fl/fl MEFs (NCLX KO) was obtained by infection with adenoviruses expressing the Cre recombinase under the control of the cytomegalovirus promoter (CMV)-Cre (ad-CRE; 300 pfu/cell; Vector Biolabs).

Mouse brain hippocampal slices were prepared from 3-month C57BL/6 mice. Briefly, animals were decapitated and brain was placed in Krebs bicarbonate dissection buffer (in mM: NaCl 120, KCl 2, CaCl2 0.5, NaHCO3 26, MgSO4 10, KH2PO4 1.18, glucose 11 and sucrose 200; pH = 7.4). After hippocampi dissection, a McIlwain Tissue Chopper was used to prepare 250 um slices. Slices were transferred to sucrose-free dissection buffer with 5 % CO_2_ for stabilization during 30 minutes before the experiment was performed.

Transfection of 30 nM siRNA or 0.25 µg S468T NCLX, adominant negative NCLX form (dnNCLX; Nita et al., 2012), kindly provided by Dr. Israel Sekler) or human NCLX (pNCLX, Adgene) was carried out using Lipofectamine 2000 (Invitrogen). Experiments were carried out 48 to 72 h after transfection.

For hypoxic treatments cells or brain slices were introduced into an Invivo2 400 workstation (Ruskinn) set at 1% O2, 5% CO2, 37 °C, and incubated for 4 or 6 hours. CGP-37158 was added 30 min before and maintained during the hypoxic treatment. N-acetyl cysteine (A7250, Sigma) or L-cysteine (30120, Sigma) were prepared in fresh before each experiment and were added two hours before hypoxic treatment and were replaced with fresh, hypoxic media immediately before hypoxic treatment.

### siRNA preparation

A double-stranded siRNA against bovine NCLX (siNCLX, sense sequence: AGCGGCCACUCAACUGCCU) was purchased from Dharmacon or Santa Cruz Biotechnology and used after proving its efficiency (Hernansanz-Agustín et al., 2020). Scrambled siRNA (siSCR) was purchased from Santa Cruz Biotechnology.

### Western blot analysis

Protein samples were extracted with reducing Laemmli buffer without bromophenol blue and quantified by Bradford assay. Extracts were loaded onto 10% standard polyacrylamide gel electrophoresis after adding 5% 2-mercaptoethanol, and subsequently transferred to nitrocellulose membranes or PVDF membranes. The following antibodies were used: polyclonal anti-NCLX antibody (ARP44042_P050, Aviva Systems Biology), mouse monoclonal anti-HIF-1α (MAB1536, R&D systems), monoclonal anti-α-tubulin antibody (#T6199, Sigma), monoclonal anti-β-actin antibody (A3854, Sigma). Antibody binding was detected by chemiluminescence with species-specific secondary antibodies labelled with horseradish peroxidase (HRP), and visualized on a digital luminescent image analyser (Fujifilm LAS-4000).

### Quantitative Real Time PCR

Total RNA was extracted using Trizol reagent (Vitro) and 0.5 µg was reverse-transcribed (Gene Amp Gold RNA PCR Core Kit; Applied Biosystems). PCR was performed with GotaqqPCR Master Mix (Promega) with 1 µL of cDNA and specific primer pairs (Table 1). β-actin mRNA was measured as an internal sample control.

**Table 1.**
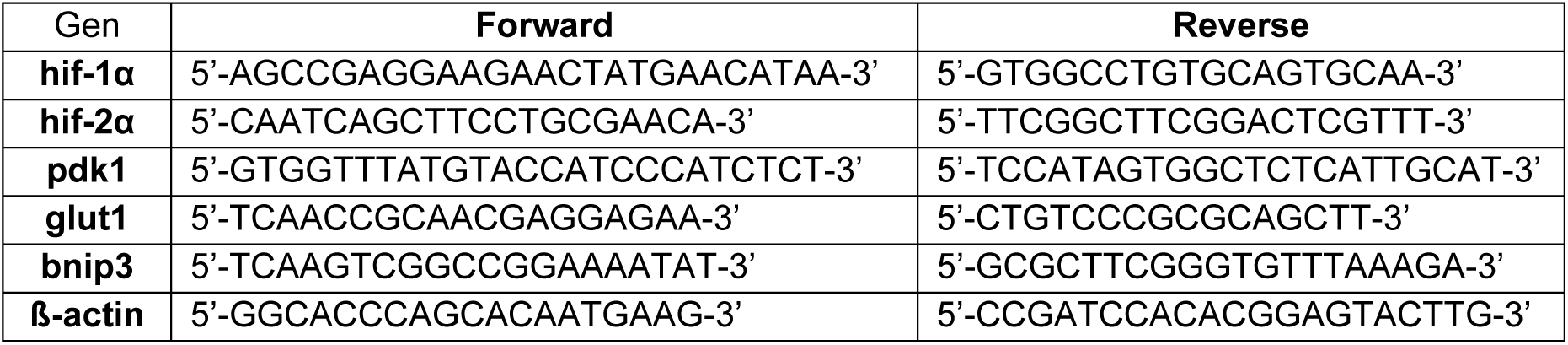
Specific primers pairs used showing the target forward and reverse sequences.

### Detection of superoxide by confocal microscopy in fixed cells

Cells were treated as previously described (Hernansanz-Agustín et al., 2021); in some conditions 10 μM CGP-37157 was added 30 min before experimentation and maintained during the experiment. Three images per each condition of 5 independent experiments were taken with a Leica SP-5 confocal microscope with a x63 objective. Excitation was performed with the 514 nm line and fluorescence emission was detected in the 560-600 nm range. Images were analysed using imageJ software as described (Hernansanz-Agustín et al., 2021).

### Cytosolic H_2_O_2_ detection with HyPer7

MAFs and MEFs were transfected with a cytosol-targeted version of Hyper7 (Pak et al., 2020). Cells were washed with PBS and maintained with HBSS + Ca/Mg. The plate was placed into a Leica SP-5 confocal microscope with an automated stage for live imaging and a thermostated cabinet, which was maintained at 37°C and 5% CO_2_. Cells were allowed to stabilize for 20 min with the new medium in the microscope stage. The objective was ×63 and excitation was performed with a 405 diode laser for the 405-nm line and an argon/krypton laser for the 488-nm line, and fluorescence emission was recorded at the 515–535-nm range. Images were quantified with Image J software.

### Statistical analysis

Data are presented as mean ± SEM. Normality and homoscedasticity tests were carried out before applying parametric tests. For comparison of multiple groups, we performed one-way analysis of variance (ANOVA) followed by Tukey’s post-hoc test. For comparison of two groups, we used Student’s two-tailed t-test. Variations were considered significant when the p value was less than 0.05. Statistical analysis and plotting were performed with GraphPad Prism 8 software.

## Results

### Hypoxic HIF-1α and HIF-2α stabilization depends on NCLX activity

Bovine aortic endothelial cells (BAECs) were exposed to four hours of either normoxia, hypoxia (1% O_2_) or the PHD inhibitor dimethyloxalglycine (DMOG), in the presence or absence of the NCLX inhibitor CGP-37157 (CGP). HIF-1α stabilization was clear upon DMOG treatment in CGP-treated or untreated cells; of note, NCLX inhibition impaired HIF-1α stabilization during hypoxia (Fig. 1a).

**Figure 1.**
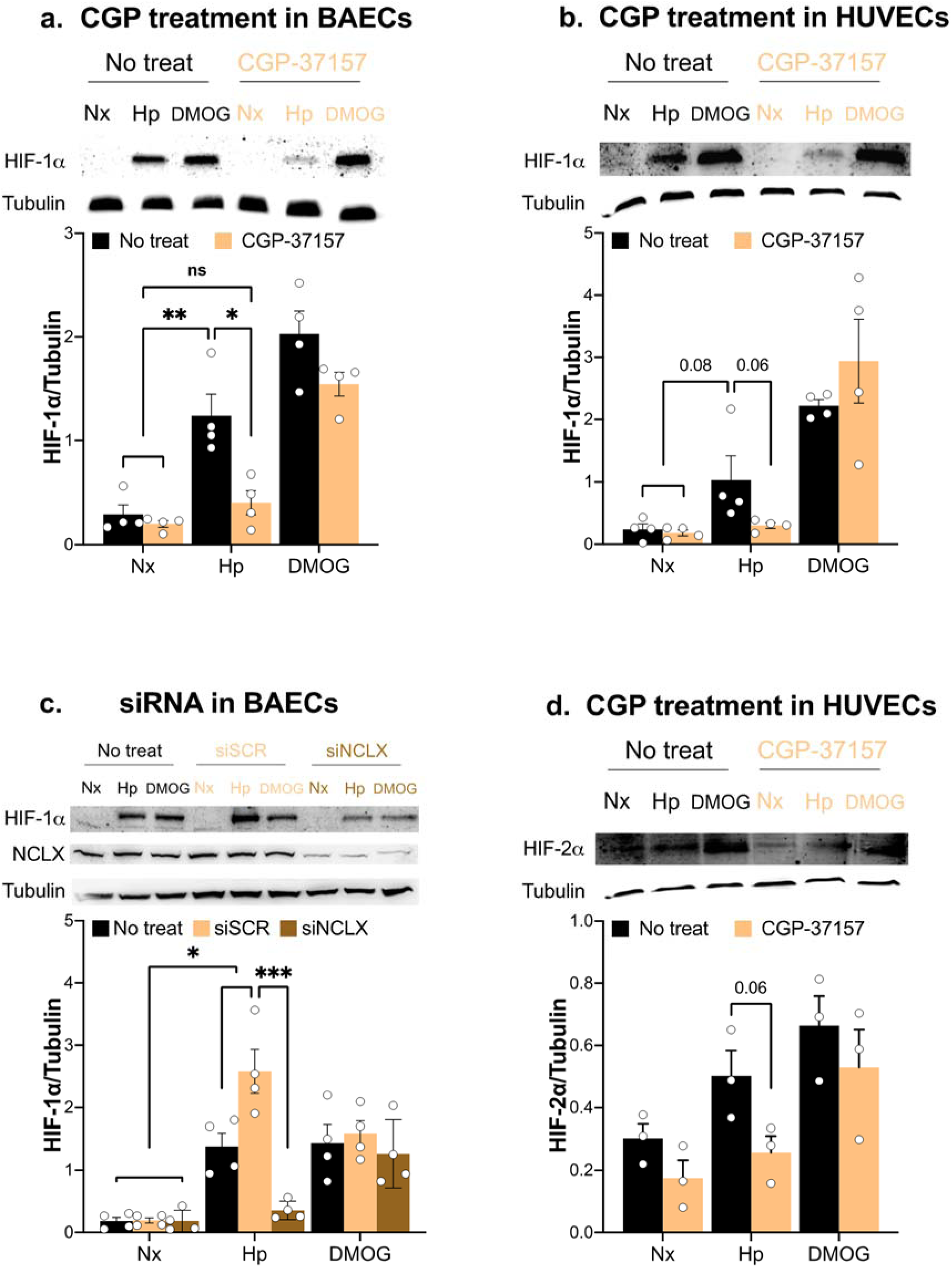
NCLX inhibition prevents HIF-α stabilization during hypoxia. HIF-1α (**a**, **b** and **c**) or HIF-2α (**d**) stabilization measured by western blotting in BAECs (**a** and **c**) or HUVECs (**b** and **d**) treated or not with 10 µM CGP-37157 (**a**, **b** and **d**) or BAECs transfected with siSCR or siNCLX (**c**). Cells were subjected for 4 hours to normoxia (Nx), normoxia with 1 mM DMOG (DMOG) or hypoxia (1% oxygen, Hp). Tubulin was used as a loading control. Representative images of at least three independent experiments are shown. *p<0.05, **p<0.01, ***p<0.001, by two-way ANOVA with Tukey’s post-hoc test.

We wondered whether human endothelial cells responded as their bovine counterparts. Thus, we used human umbilical vein endothelial cells (HUVECs) under the same experimental rationale. HUVECs exposed to DMOG stabilized HIF-1α in CGP-treated and untreated cells. Again, HIF-1α stabilization was greatly impaired during hypoxia after NCLX inhibition (Fig. 1b).

To test the specificity of the treatment, we silenced NCLX expression with a specific siRNA for its gene, *Slc8b1*, which was effective diminishing NCLX protein levels compared to the treatment with control scramble siRNA (Fig. 1c). Neither treatment was able to inhibit DMOG-induced HIF-1α stabilization; however, NCLX silencing blocked HIF-1α stabilization under hypoxia (Fig. 1c).

We also assessed HIF-2α stabilization in CGP-treated or untreated HUVECs exposed to either four hours of normoxia, hypoxia or DMOG. As expected, DMOG induced the stabilization of HIF-2α in either condition. On the contrary, HIF-2α stabilization in CGP-treated HUVECs was clearly diminished with respect to untreated cells (Fig. 1d), showing a similar behaviour as for HIF-1α.

### HIF-1-dependent transcriptional activity depends on NCLX activity

In order to test the activity of HIF-1 in response to hypoxia, assessing the response to hypoxia through the HIF pathway, we performed RT-PCR of known HIF-1 target genes (Althaus et al., 2006; Kim et al., 2006; Ouiddir et al., 1999). HUVECs were exposed to six hours of either normoxia, hypoxia or DMOG in the presence or absence of CGP. We first tested whether CGP treatment under any of the conditions modified the levels of hif-1α or hif-2α mRNAs. Neither hif-1α or hif-2α mRNA levels were affected by NCLX inhibition (Fig. 2a-b).

**Figure 2.**
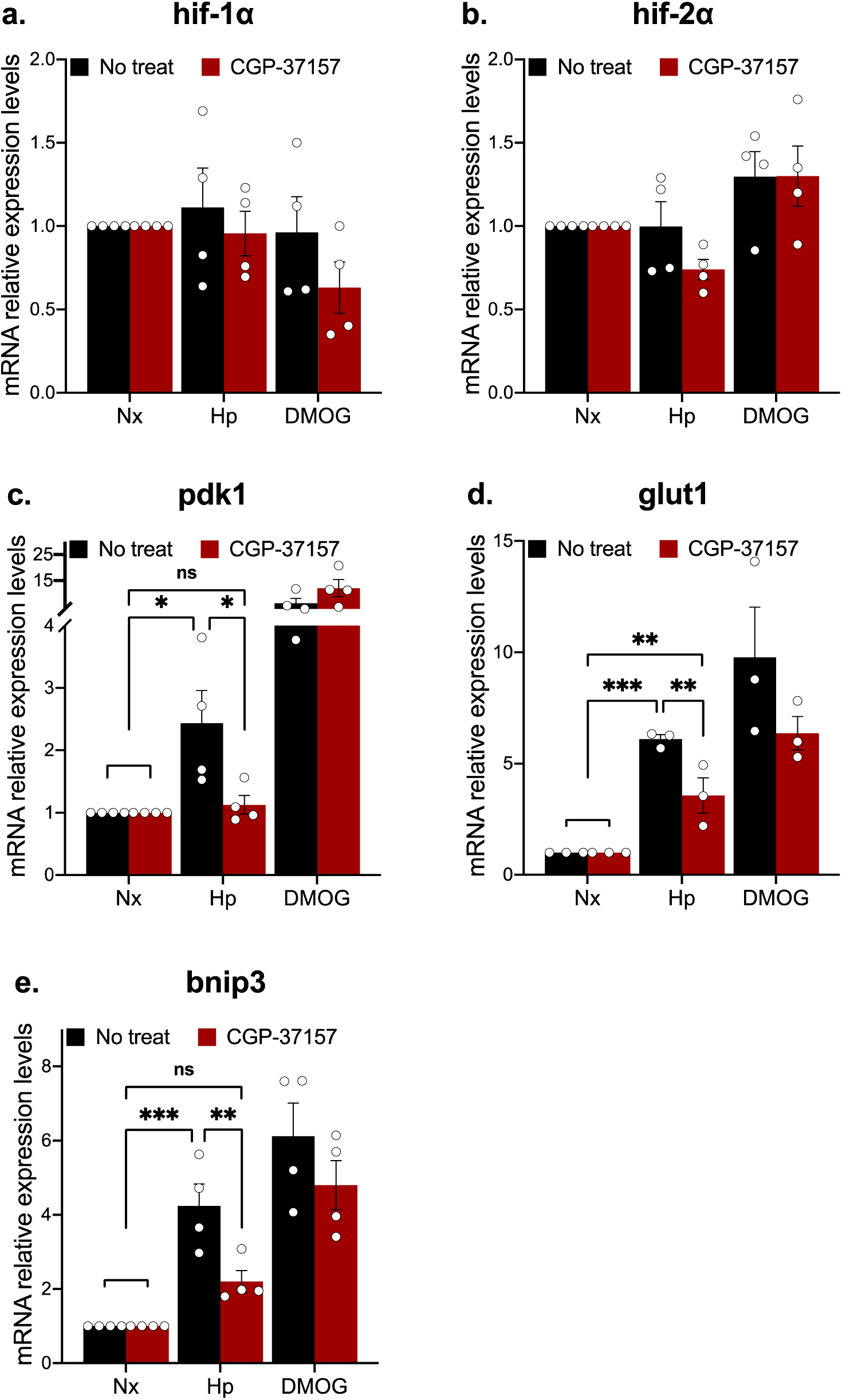
NCLX inhibition prevents HIF pathway activation. Quantitative RT-PCR analysis of hif-1α (**a**), hif-2α (**b**), pdk1 (**c**), glut1 (**d**) and bnip3 (**d**) mRNAs in untreated (black) or 10 µM CGP-37157-treated (red) HUVECs subjected to 6 h of normoxia (Nx), hypoxia (1% oxygen, Hp) or normoxia with 1 mM DMOG (DMOG). *p<0.05, **p<0.01, ***p<0.001, by two-way ANOVA with Tukey’s post-hoc test.

Pyruvate dehydrogenase kinase 1 (PDK1), an enzyme controlling the fate of pyruvate into the mitochondria, prominently increased its expression upon DMOG treatment in either condition, indicating a clear activation of the HIF-1 pathway. However, hypoxia only increased the levels of pdk1 in untreated cells, in contrast to NCLX-inhibited cells, indicating that NCLX was involved in HIF-1α stabilization (Fig. 2c).

The glucose transporter Glut1 is a HIF-1 target that facilitates the entrance of glucose into the cell, in this way being involved in the hypoxic glycolytic switch. DMOG treatment promoted glut1 expression in the presence or absence of CGP. However, the effect of hypoxia was clearly diminished in NCLX-inhibited cells with respect to the increase observed in untreated cells (Fig. 2d).

We selected a third HIF-1 target, the BCL2 interacting protein 3 (BNIP3). Whereas DMOG treatment showed a clear increase of bnip3 expression in both untreated and CGP-treated cells, NCLX inhibition clearly affected the increase induced by hypoxia treatment (Fig. 2e).

So far, these data indicate that NCLX activity is necessary for the stabilization of HIF-1α and HIF-2α, and for the activation of HIF-1 during hypoxia, but not under PHD inhibition by DMOG.

### NCLX inhibition prevents hypoxic HIF-1α stabilization in a neuronal cell line and hippocampus

We next wondered whether more oxidative cells, such as the neuronal cell line HT-22, were also sensitive to NCLX inhibition regarding HIF-1α stabilization. HT-22 cells were subjected to four hours of normoxia, hypoxia or DMOG. DMOG treatment yielded a strong stabilization of HIF-1α in both CGP-treated and untreated cells. However, only untreated cells were able to stabilize HIF-1α during hypoxia (Fig. 3a).

**Figure 3.**
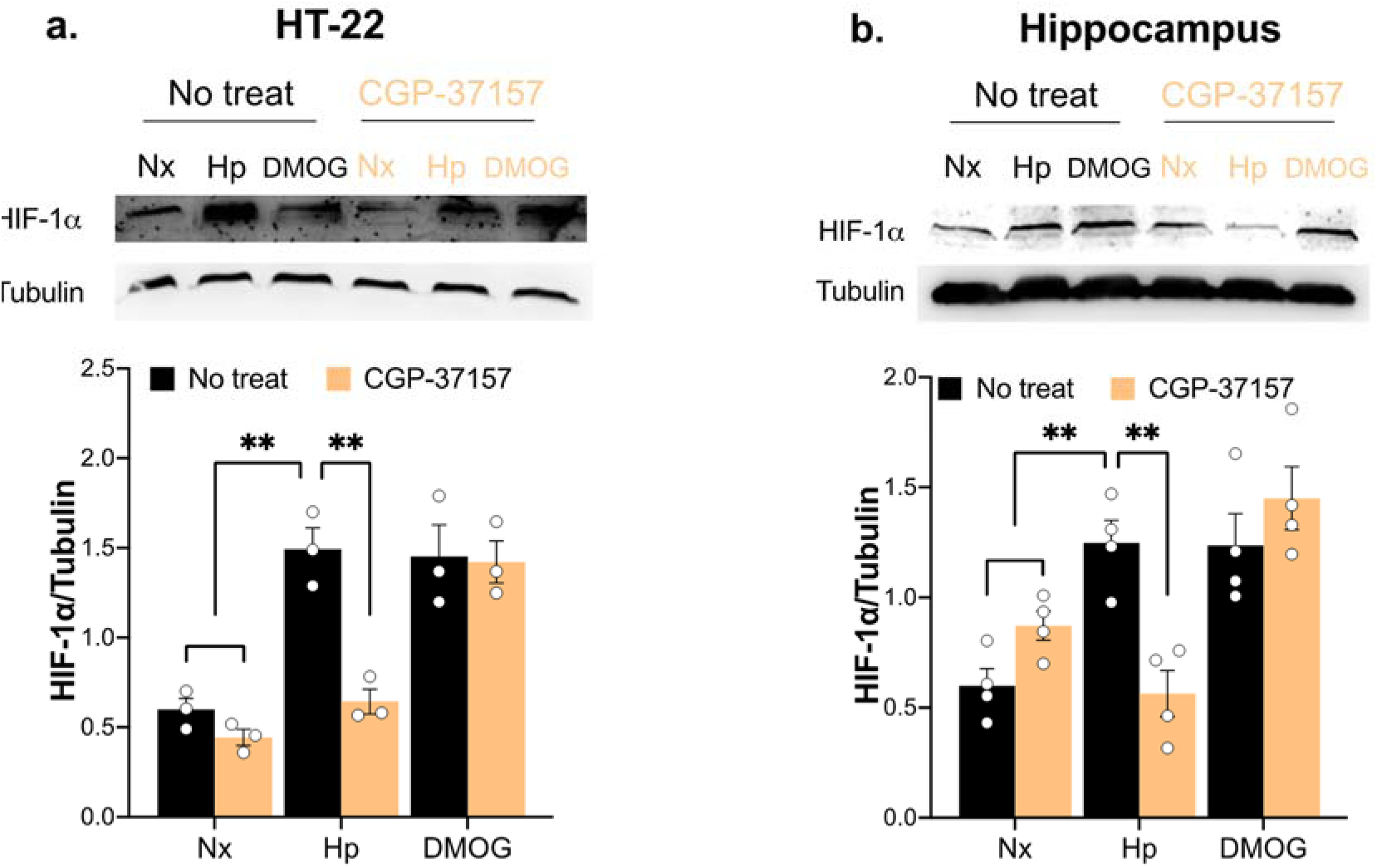
NCLX inhibition abolishes hypoxic HIF-1α stabilization in neuronal cell line and hippocampus. HIF-1α stabilization measured by western blotting in HT-22 cells (**a**) or hippocampus slices (**b**) treated or not with 10 µM CGP-37157. Cells or slices were subjected for 4 hours to normoxia (Nx), normoxia with 1 mM DMOG (DMOG) or hypoxia (1% oxygen, Hp). Tubulin was used as a loading control. Representative images of at least three independent experiments are shown. **p<0.01, by two-way ANOVA with Tukey’s post-hoc test.

As neuronal cell lines undergo adaptation to cell culture and may become more glycolytic than a primary neuronal culture, we tested the same hypothesis on hippocampal tissue. We extracted hippocampal slices from mice and subjected them to either four hours of normoxia, hypoxia or DMOG. As with the neuronal cell line, DMOG induced a clear stabilization of HIF-1α; however, hypoxic slices were only able to stabilize HIF-1α when NCLX was active (Fig. 3b).

### NCLX inhibition does not prevent HIF-1α stabilization during hypoxia in mouse embryonic fibroblasts, unless they are pre-treated with N-acetyl-L-cysteine (NAC) or L-cysteine

We wondered whether chronic NCLX impairment also prevented the stabilization of HIF-1α during hypoxia in mouse embryonic fibroblasts (MEFs). Wild type (WT) and *Slc8b1* knock-out (KO) MEFs were subjected to four hours of either normoxia, hypoxia or DMOG. In both cell types, DMOG was able to stabilize HIF-1α. Surprisingly, genetic abolition of NCLX did not prevent the stabilization of HIF-1α during hypoxia (Fig. 4a), though they were unable to produce hypoxic ROS (Fig. 4b). Thus, we tested whether acute NCLX inhibition affected HIF-1α stabilization in MEFs. Neither pharmacological NCLX inhibition using CGP (Fig. 4c) nor overexpression of a dominant negative form of NCLX (dnNCLX) (Fig. 4d) prevented hypoxic HIF-1α stabilization in MEFs. We then assessed if the response of MEFs to hypoxia might differ from other cell types, in which the superoxide burst produced during the first minutes of hypoxia depends on NCLX activity (Hernansanz-Agustín et al., 2020). However, we observed that MEFs produce a superoxide burst in response to acute hypoxia when NCLX is active, but not when it is pharmacologically inhibited with CGP (Fig. 4b). Thus, it seems that HIF-1α stabilization after 4 hours of hypoxia does not depend on NCLX activity or acute hypoxic ROS production in this cell type under these experimental conditions.

**Figure 4.**
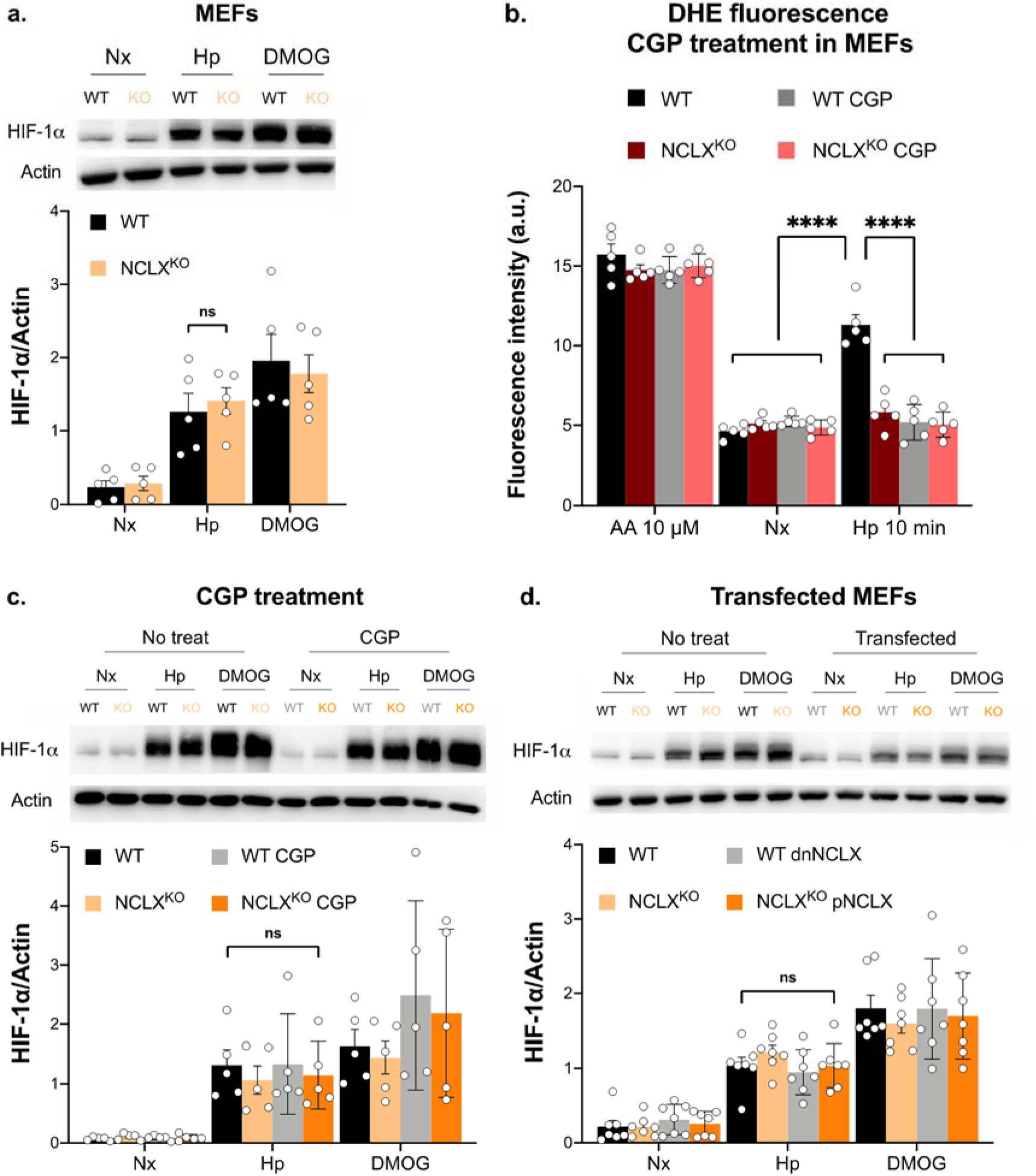
NCLX inhibition does not prevent HIF-1α stabilization during hypoxia in MEFs. (**a**, **c** and **d**) HIF-1α stabilization measured by western blotting in MEFs. WT and NCLX KO MEFs untreated (**a**) or treated with 10 µM CGP (**c**) or transfected with dnNCLX or pNCLX (**d**) were exposed for 4 hours to normoxia (Nx), normoxia with 1 mM DMOG (DMOG) or hypoxia (1% oxygen, Hp). ß-actin was used as a loading control. Representative images of at least five independent experiments are shown. (**b**) Detection of superoxide production by fluorescence microscopy in fixed cells. In normoxia, MEFs were incubated for 30 min with antimycin A (AA 10 µM) or without AA (Nx) and some cells were treated with 10 µM CGP; then, DHE was added for 10 min and cells were fixed. For hypoxic condition, some cells were treated with 10 µM CGP for 30 min in normoxia and then, were transferred to hypoxia chamber (1% oxygen). Inside it, medium was replaced for pre-hypoxic-balanced medium and, quickly, DHE was added for 10 min and cells were fixed inside hypoxia chamber. ****p<0.0001, by two-way ANOVA with Tukey’s post-hoc test.

We previously noticed that basal H_2_O_2_ levels, after NCLX knock-out in MEFs, were higher than in WT MEFs (Hernansanz-Agustín et al., 2020). To discern whether higher basal H_2_O_2_ had a role in HIF-1α stabilization during hypoxia we treated both NCLX WT and KO MEFs with 1 mM NAC for two hours during normoxia (NAC pre-treatment) and then cells were transferred into the hypoxic chamber, inside which the media was replaced with fresh, hypoxic-stabilized media without NAC. Again, WT and KO MEFs without NAC pre-treatment stabilized HIF-1α during hypoxia (Fig. 5a). NCLX WT MEFs pre-treated with NAC were able to stabilize HIF-1α; however, NAC-pre-treated KO MEFs were unable to stabilize HIF-1α during hypoxia (Fig. 5a). Similar results were obtained with L-cysteine addition: only L-cysteine pre-treatment prevented HIF-1α stabilization during hypoxia in NCLX KO MEFs (Fig. 5b).

**Figure 5.**
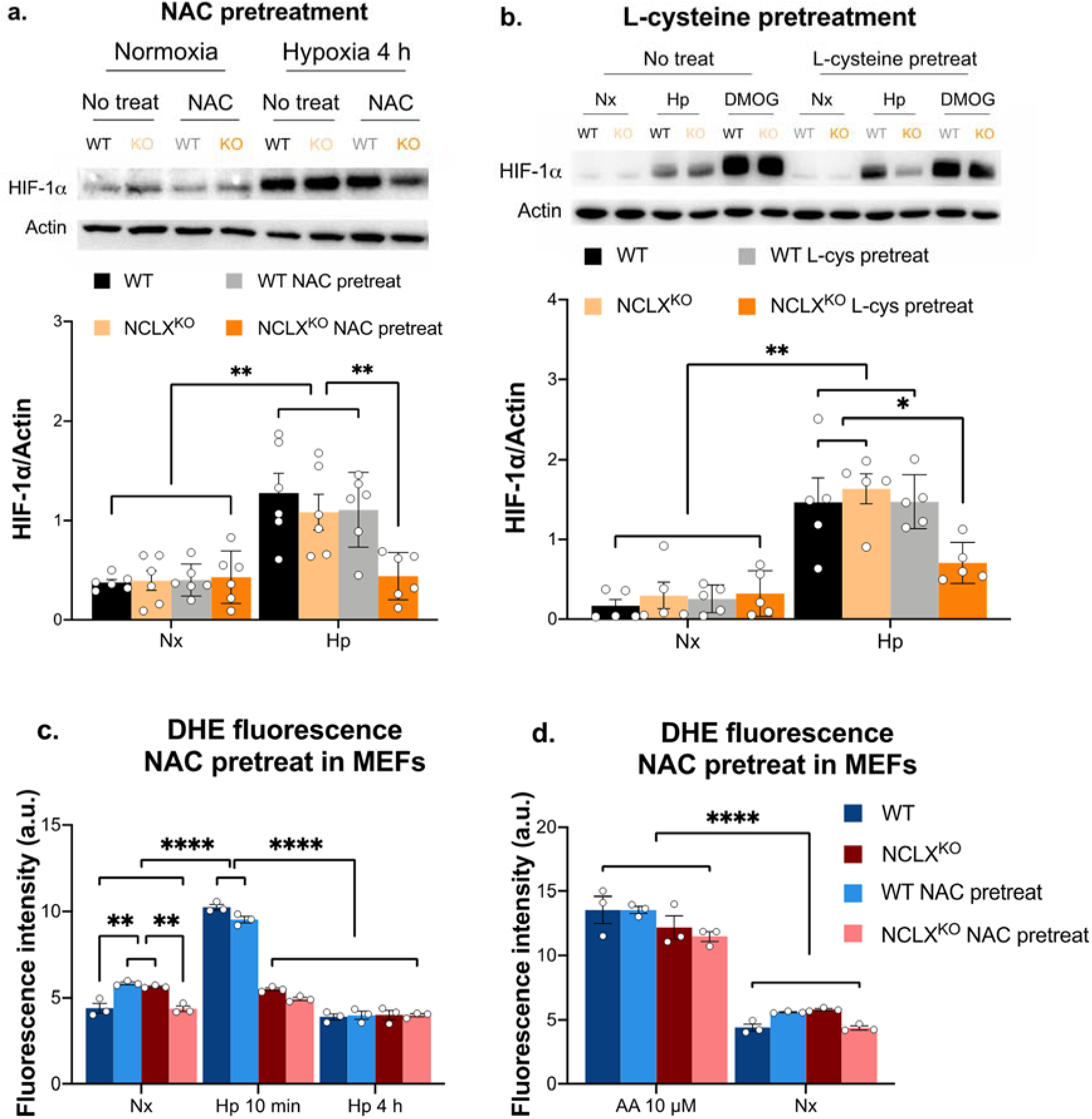
Effects of NAC pre-treatment on HIF-1α stabilization and superoxide production during hypoxia in MEFs. (**a** and **b**) HIF-1α stabilization measured by western blotting in MEFs. WT and NCLX KO MEFs were treated with 1 mM NAC (**a**) or 1 mM L-cysteine (**b**) for 2 hours in normoxia. Then, cells were incubated with fresh media in normoxia for 4 hours (Nx) or with pre-hypoxic-balanced media in the hypoxia chamber (1% oxygen) for 4 hours (Hp). ß-actin was used as loading control. Representative images of at least five independent experiments are shown. (**c** and **d**) Detection of superoxide production by fluorescence microscopy in fixed cells. Some cells were treated for 2 hours with 1 mM NAC in normoxia. For normoxia conditions (**c** and **d**), cells were incubated with fresh media in normoxia for 4 hours (Nx) or, during the last 30 min of these 4 hours, some cells were treated with 10 µM antimycin A (AA). Then, DHE was added for 10 min and cells were fixed. For hypoxia conditions (**c**), medium with NAC treatment was replaced by pre-hypoxic-balanced medium in the hypoxia chamber (1% oxygen) for 10 min (Hp 10 min) or for 4 hours (Hp 4 h). Then, DHE was added for 10 min and cells were fixed inside the hypoxia chamber. At least three independent experiments were done. *p<0.05, **p<0.01, ****p<0.0001, by two-way ANOVA with Tukey’s post-hoc test.

Using DHE, we confirmed that basal superoxide levels are slightly increased in NCLX KO MEFs compared to WT (Fig. 5c). Nevertheless, while WT MEFs pre-treated for 2 hours with NAC before the 4 hours in hypoxia showed an increase in ROS production, NCLX KO MEFs pre-treated with NAC showed a decrease in ROS production (Fig. 5c). However, NAC pretreatment did not alter the superoxide burst in the first minutes of hypoxia, which was maintained in WT MEFs but not in KO MEFs (Fig. 5c). Antimycin A-induced superoxide production was neither affected by NAC pretreatment (Fig. 5d).

These results point out that the pre-treatment of MEFs with NAC or L-Cys restores the dependency of NCLX activity for hypoxic HIF-1α stabilization observed in many other cell types. We wondered what the reason was behind such special behaviour of MEFs in comparison to many other cell types.

### NCLX inhibition prevents hypoxic HIF-1α stabilization in mouse adult fibroblasts (MAFs)

Since MEFs are embryonic cells, we decided to check HIF-1α stabilization in isogenic mouse adult fibroblasts. MAFs were subjected to four hours of normoxia, hypoxia or DMOG with or without NCLX inhibition. DMOG treatment induced the stabilization of HIF-1α in both CGP-treated and untreated cells, while HIF-1α stabilization during hypoxia was clearly impaired after NCLX inhibition with CGP-37157 (Fig. 6a). Similar results were obtained in MAFs overexpressing dNCLX, which also impaired HIF-1α stabilization in hypoxia (Fig. 6b). We also wondered if the hypoxic superoxide burst in MAFs was NCLX-dependent. We observed that MAFs produce a superoxide burst in response to acute hypoxia when NCLX was active, but not when it was pharmacologically blocked with CGP (Fig. 6c). Hence, in adult fibroblasts NCLX-dependent hypoxic-ROS production is linked to HIF-1α stabilization.

**Figure 6.**
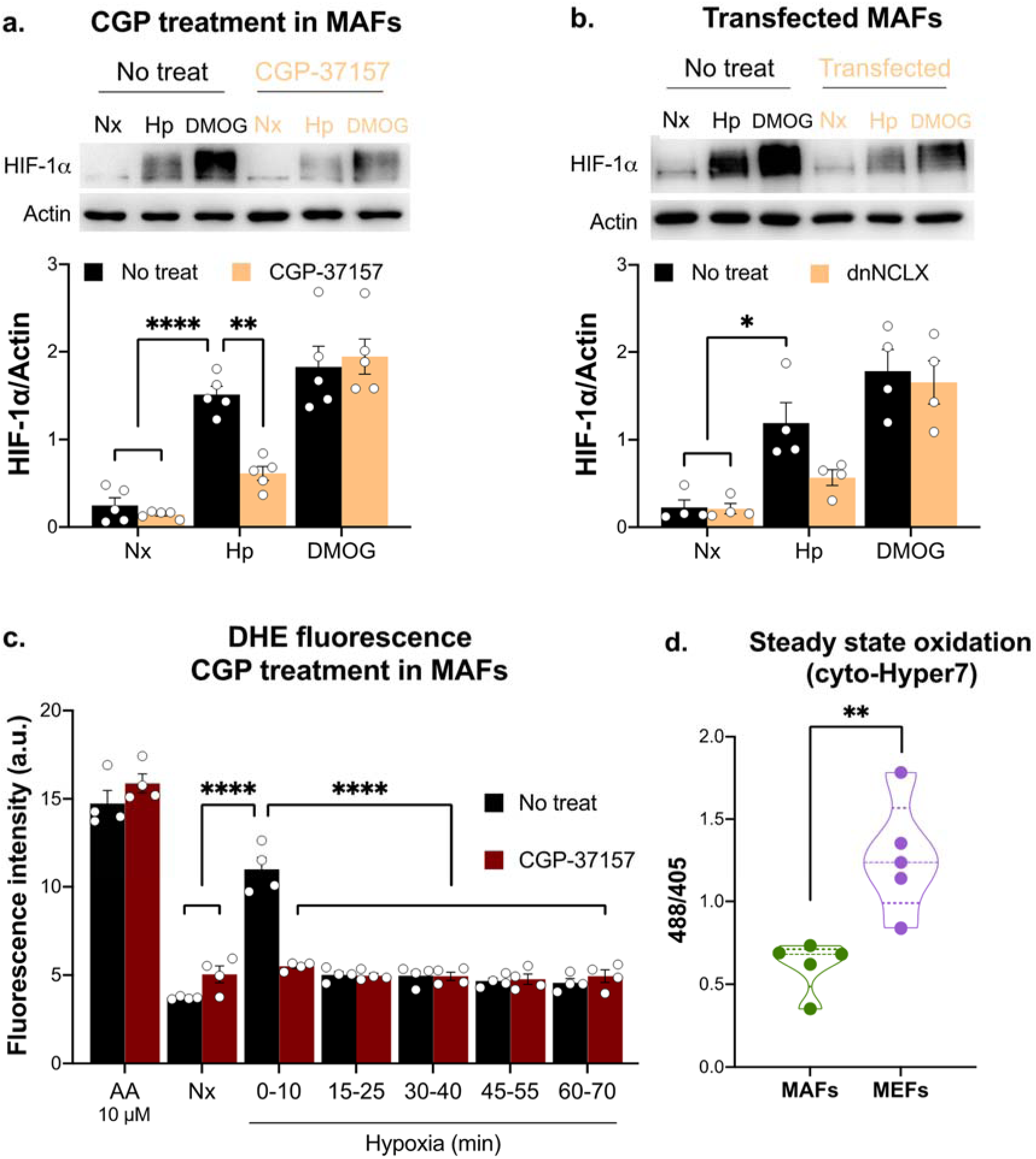
NCLX inhibition prevents HIF-1α stabilization during hypoxia in MAFs. (**a** and **b**) HIF-1α stabilization measured by western blotting in MAFs. MAFs treated with 10 µM CGP (**a**) or transfected with dnNCLX (**b**) were exposed for 4 hours to normoxia (Nx), normoxia with 1 mM DMOG (DMOG) or hypoxia (1% oxygen, Hp). ß-actin was used as loading control. Representative images of at least four independent experiments are shown. (**c**) Detection of superoxide production by fluorescence microscopy in fixed cells. For normoxic conditions, MAFs were incubated for 30 min with antimycin A (AA 10 µM) or without AA (Nx) and some cells were treated with 10 µM CGP. Then, DHE was added for 10 min and cells were fixed. For hypoxic conditions, some cells were treated with 10 µM CGP for 30 min in normoxia prior to hypoxia and, then, were transferred to hypoxia chamber (1% oxygen). Inside it, medium was replaced for pre-hypoxic-balanced medium for 0, 15, 30, 45 or 60 min and then DHE was added for 10 min more and cells were fixed inside hypoxia chamber. (**d**) Detection of cytosolic basal ROS levels by a cytosolic version of Hyper7 with microscopy. At least four independent experiments were done. *p<0.05, **p<0.005, ****p<0.0001, by two-way ANOVA with Tukey’s post-hoc test (**a-c**) or Student’s t test (**d**).

It is known that the amount and activity of some antioxidant proteins depend on development (Surai et al., 2016), so we wondered whether basal ROS production could be different in MEFs than in MAFs. Thus, we used the redox-sensitive protein cyto-HyPer7 (Pak et al., 2020). We observed that the steady state of cytosolic ROS levels is higher in MEFs than in MAFs (Fig 6d), suggesting that the hypoxic redox target leading to HIF-1α stabilization may be already oxidized in MEFs, producing that the medium-term adaptation to hypoxia, in this specific cell type, is independent of hypoxic mitochondrial ROS production.

## Discussion

Human and other mammalian organisms adapt to hypoxia by a great variety of metabolic, cellular and physiological responses, which differ in the times of response and the molecular mechanisms implied in oxygen sensing and signal transduction. The precise oxygen sensor for acute responses to hypoxia has not yet been identified, although several molecular mediators have been described. Among these mediators, changes in the NADH/NAD^+^ redox ratio and increased ROS production have been identified, in which mitochondrial functions are implied (Brunelle et al., 2005; Chandel et al., 1998; Guzy et al., 2005; Hernansanz-Agustín et al., 2017; Kim et al., 2006; Mansfield et al., 2005). On the contrary, the oxygen sensor underlying hypoxia adaptation through regulation of gene expression mediated by HIFs was characterized two decades ago, and the three leading investigators of this mechanism were awarded the Nobel Prize in Physiology or Medicine in 2019 (Kaelin & Ratcliffe, 2008).

Even though the PHDs, the oxygen sensors for the HIF pathway, are located in the cytosol and the nucleus, a role of mitochondria on HIF-1α stabilization in hypoxia has been debated since HIF-1α was identified, before and after the identification of the PHDs (Chandel et al., 1998; Guzy & Schumacker, 2006; Hagen et al., 2003; Kaelin & Ratcliffe, 2008). Most of the approaches studying the mitochondrial function tried to block ROS production, which involved the inhibition or ablation of the mitochondrial respiratory chain and the alteration of the cellular O_2_ gradients and metabolism, which can be confusing due to the alteration of the several functions that are overlapped in the mitochondrial electron transport chain complexes. On the other hand, defects in mitochondrial proteins not belonging to the electron transport chain or the Krebs have been shown to affect HIF-1α stabilization in different ways, not directly correlated with changes in ROS levels (Shvetsova et al., 2017)

The mitochondrial Na^+^/Ca^2+^ exchanger was identified time ago as the NCLX protein (Palty et al., 2010). We have recently shown that NCLX activity is necessary for producing a superoxide burst in the first minutes of hypoxia, which is involved in acute responses such as pulmonary vasoconstriction (Hernansanz-Agustín et al., 2020). We now show that NCLX inhibition, which blocks hypoxic ROS production by mitochondria without affecting mitochondrial respiration (Hernansanz-Agustín et al., 2020), inhibited the hypoxic HIF-1α and HIF-2α stabilization in different cell types and in hippocampal slices. This highlights the role of NCLX activity, and most probably of mitochondrial Na^+^ import, in regulating responses to hypoxia mediated by the activation of the HIF pathway. In line with this, the elevation of intracellular Na^+^ has recently been shown to produce a major metabolic reprogramming of cardiac metabolism through NCLX activity (Aksentijević et al., 2020), including an increase in glycolytic activity that is a signature of HIF-driven metabolic adaptation (Semenza, 2013).

However, our results also show that a particular cell type, MEFs, were able to stabilize HIF-1α in response to hypoxia with or without functional NCLX. It is important to highlight that MEFs have been the only embryonic cells used in this study, and these cells show higher basal ROS levels comparied with their correspondent adult cells, MAFs (Fig. 6d). In addition, in MEFs the dependency of HIF-1α stabilization upon NCLX activity is restored after pre-treatment with NAC or L-cysteine (Fig. 5a and 5b), both well-known intracellular antioxidant defence enhancers. In line with this, in a colorectal cancer mouse model (Pathak et al., 2020), chronic NCLX deletion induced HIF-1α stabilization without hypoxic stimulus and it was prevented by adding mito-TEMPO, a mitochondria-targeted ROS scavenger. Antioxidant-enhancing capacity of NAC and L-cysteine may decrease higher basal H_2_O_2_ levels in WT and KO MEFs, reversing a rather pro-oxidant cellular environment which made MEFs insensitive to hypoxic NCLX-dependent H_2_O_2_ production. After NAC or L-cysteine pre-treatment the dependency of HIF-1α stabilization on NCLX is restored, resembling other cell types in this study. Taken together, these results suggest that HIF-1α stabilization during hypoxia is dependent on hypoxic ROS production according to the redox cellular context: if the basal ROS levels are *per se* elevated, NCLX-dependent hypoxic ROS are not necessary to induce HIF-1α stabilization during hypoxia; however, if the basal ROS levels are relatively low, then NCLX-dependent hypoxic ROS must occur to induce HIF-1α stabilization during hypoxia (Figure 7).

**Figure 7.**
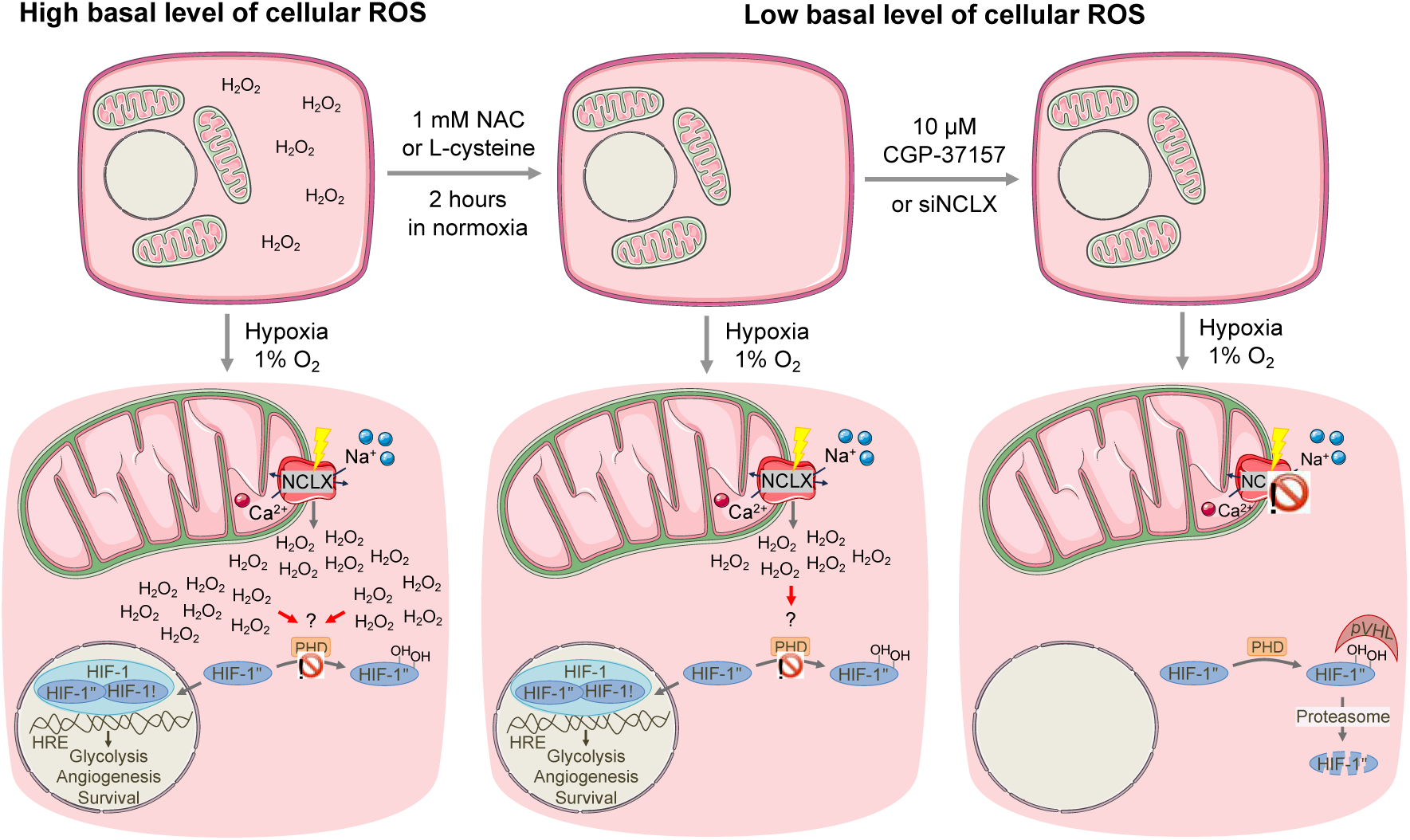
NCLX-dependent hypoxic ROS are needed for HIF-1α stabilization depending on cellular ROS context. Cells with high basal level of ROS (such as MEFs) subjected to hypoxia (1% oxygen) present enough cytosolic ROS to inactivate the prolyl hydroxylases (PHDs) in hypoxia and induce HIF-1α stabilization; meaning that the hypoxic NCLX-dependent ROS production is unnecessary for HIF-1α stabilization. After its stabilization, HIF-1α together with β subunit are bound to the hypoxia response elements (HREs), inducing the genetic responses related to glycolysis and angiogenesis, among others. In cells with low basal level of ROS (such as MAFs) or after decreasing basal level of ROS, by treating cells with 1 mM NAC or 1 mM L-cysteine for 2 hours in normoxia prior to hypoxia, the hypoxic NCLX-dependent ROS production is recovered as the source of ROS for PHDs inactivation and, in turn, HIF-1α stabilization occurs. Therefore, by blocking NCLX with 10 µM CGP-37157 or with a siRNA against NCLX (siNCLX), hypoxic ROS production is also inhibited and, thus, PHDs are active and hydroxylate HIF-1α. The Von Hippel-Lindau tumor suppressor (pVHL) recognizes HIF-1α hydroxylated which, eventually, tags HIF-1α to be degraded through proteasome and, thus, hypoxic HIF-1α stabilization is avoided when NCLX is inhibited. Question mark means that the ROS-dependent PHD inhibition may be mediated by a redox target and prohibition sign means protein activity inhibition. Scheme made using illustrations available on Elsevier Medical Art

Additionally, intracellular cysteine level has been shown to work as a PHD regulator (Briggs et al., 2016). The authors postulated that specific intramolecular cysteine residues of PHD are prone to oxidation, inducing an auto-inhibition of PHD, so that treatment with free cysteine reverses that oxidation, either directly or through increasing reduced glutathione (GSH), which has also been shown to increase PHD activity (Nytko et al., 2011). However, we have not identified if this effect on ROS levels is directly impacting on PHD or if it is due to some redox-sensitive transductor.

Nevertheless, NAC or L-cysteine pre-treatment may be inducing ROS-independent effects which can be participating in this mechanism. For instance, the intracellular cysteine depletion observed in (Briggs et al., 2016) was a consequence of the excessive glutamate export by cells, which in turn inhibited their own xCT glutamate-cystine antiporter. Consequently, pre-treatment with NAC or L-cysteine might be altering some metabolic pathways in MEFs, thereby reinstating the NCLX-dependency of HIF-1α stabilization.

Our observations highlight the involvement of NCLX in the stabilization of HIF-1α during hypoxia in different cell types. Our previous work described a role for NCLX in triggering responses to acute hypoxia. The current findings provide an additional role of NCLX in the responses to hypoxia in a longer timeframe that are driven by changes in gene expression. Further studies would be needed to address if these two effects use similar or different molecular mechanisms as well as the implication of NCLX activity in different physiological and pathological responses to hypoxia.

## Acknowledgements

We thank Tamara Villa-Piña, Elena Ramos, Tamara Oliva and Esther Fuertes-Yebra for technical assistance. We also thank Dr. John Elrod (Temple University, USA), Dr. José Antonio Enríquez (CNIC, Spain) and Dr. Israel Sekler (Ben Gurion University of the Negev, Israel) for providing us materials needed to complete the study.

This research has been financed by grants from the Spanish Government (partially funded by the European Union ERDF “A way of making Europe” and NextGenerationEU): RedoxStroke (RTI2018-094203-B-I00), excellence network RED2018-102576-T, NCLRedoX (PID2021-124688OB-I00), NCLXtroke (PDC2022-133246-I00), PID2021-125986OB-I00 and PID2022-139936OA-I00 from AEI (MICIU/AEI/10.13039/501100011033); PI15/00107 , PI22/00362 and RICORS-ICTUS (RD21/0006/0009) from Instituto de Salud Carlos III (ISCIII) and fellowship FPU18/03475 to C.C.-F. from MICIU, and by a grant from the Fundación Domingo Martínez.

